# TopDIA: A Software Tool for Top-Down Data-Independent Acquisition Proteomics

**DOI:** 10.1101/2024.04.05.588302

**Authors:** Abdul Rehman Basharat, Xingzhao Xiong, Tian Xu, Yong Zang, Liangliang Sun, Xiaowen Liu

**Author notes:** Correspondence: Xiaowen Liu; Liangliang Sun. Co-first authors (Abdul Rehman Basharat, Xingzhao Xiong).

## Abstract

Top-down mass spectrometry is widely used for proteoform identification, characterization, and quantification owing to its ability to analyze intact proteoforms. In the last decade, top-down proteomics has been dominated by top-down data-dependent acquisition mass spectrometry (TD-DDA-MS), and top-down data-independent acquisition mass spectrometry (TD-DIA-MS) has not been well studied. While TD-DIA-MS produces complex multiplexed tandem mass spectrometry (MS/MS) spectra, which are challenging to confidently identify, it selects more precursor ions for MS/MS analysis and has the potential to increase proteoform identifications compared with TD-DDA-MS. Here we present TopDIA, the first software tool for proteoform identification by TD-DIA-MS. It generates demultiplexed pseudo MS/MS spectra from TD-DIA-MS data and then searches the pseudo MS/MS spectra against a protein sequence database for proteoform identification. We compared the performance of TD-DDA-MS and TD-DIA-MS using *Escherichia coli* K-12 MG1655 cells and demonstrated that TD-DIA-MS with TopDIA increased proteoform and protein identifications compared with TD-DDA-MS.

## Introduction

Top-down mass spectrometry (MS) has become the method of choice for identifying and quantifying intact proteoforms^1, 2^ in biological samples as it provides a bird’s-eye view of entire proteoforms. Recent advances in proteoform separation techniques and MS have significantly enhanced proteoform identifications in human cells and samples using top-down proteomics (TDP) ^3-5^, thus enabling proteoform profiling and the identification of differentially expressed proteoforms.

Most TDP studies use top-down data-dependent acquisition MS (TD-DDA-MS) for the selection of proteoform ions for tandem mass spectrometry (MS/MS) analysis^6, 7^. In TD-DDA-MS, the mass spectrometer performs a sequential survey of all precursor ions as they elute from the separation system, such as liquid chromatography (LC)^8, 9^ or capillary electrophoresis (CE)^10, 11^, and the *N* most intense ions are isolated in each MS1 scan and fragmented to produce MS/MS spectra^12, 13^. These MS/MS spectra are utilized to identify proteoforms by protein sequence database search. However, the reproducibility of proteoform identifications is limited due to the stochastic nature of the precursor ion selection in TD-DDA-MS experiments^6, 14-16^.

Unlike TD-DDA-MS, which focuses on acquiring MS/MS data for top-intensity precursor ions, top-down data-independent acquisition MS (TD-DIA-MS) generates fragments for all precursor ions within preselected isolation windows^13, 17^, resulting in an LC-MS/MS map for each isolation window. By collecting MS/MS spectra from all precursor ions, TD-DIA-MS provides comprehensive MS/MS data acquisition without the need for prior precursor information^17-19^. Consequently, TD-DIA-MS has the potential to circumvent certain shortcomings of TD-DDA-MS, including its low reproducibility and tendency to fail to generate MS/MS spectra of low-intensity precursor ions. However, TD-DIA-MS also presents its challenges, notably the complexity of multiplexed MS/MS spectra generated by the co-fragmentation of multiple precursor ions.

Many methods have been proposed for identifying peptides from bottom-up^20^ DIA-MS data. These methods can be categorized into two classes: spectral library-based methods and library-free methods^21, 22^. Spectral library-based tools such as OpenSWATH^23^, Spectronaut^14^, Skyline^24^, Specter^25^, and EncyclopeDIA^26^ rely on spectral libraries for peptide identification from DIA-MS data. These libraries are created through a database search of DDA-MS or DIA-MS data from the same sample^27, 28^, or deep-learning models, such as DeepMass^29^, pDeep^30^, Prosit^31^, and DeepDIA^32^. Library-free tools, such as DIA-Umpire^33^, Group-DIA^34^, directDIA^14^, Dear-DIA^XMBD35,^ and PECAN^36^, rely on protein sequence databases, not spectral libraries for peptide identification^37, 38^. Some tools, such as DIA-NN^39^, and MaxDIA^40^, offer support for both library-based and library-free approaches.

In TDP, the analysis of intact proteoforms^41^ results in ions with large masses and high charge states. In general, top-down mass spectra of intact proteoforms are more complex than bottom-up mass spectra of digested peptides^41-44^. Consequently, computational methods designed for bottom-up DIA-MS data analysis are often inefficient for processing TD-DIA-MS data. Here, we present TopDIA, the first spectrum-centric^37^ software tool for proteoform identification using TD-DIA-MS. TopDIA generates pseudo non-multiplexed MS/MS spectra from TD-DIA-MS data and searches these pseudo spectra against a protein sequence database for proteoform identification. Experimental results demonstrate that TD-DIA-MS with TopDIA identified 9.3% more proteoforms and 10.5% more proteins from *Escherichia coli (E. coli)* K-12 MG1655 cells compared with TD-DDA-MS.

## Methods

### Sample preparation

*E. coli* K-12 MG1655 cells were inoculated in 2 mL sterilized LB medium at 37 °C with shaking at 250 rpm for 2 hours. Subsequently, a 250 mL *E. coli* sample was transferred into 400 mL LB solution and incubated with shaking at 250 rpm at 37 °C for 16 hours. The solution was centrifuged at 5,000 g for 5 min at 4 °C. Cell pellets were collected, washed using 5 mL 1x phosphate-buffered saline (PBS) three times, and then resuspended in 200 uL 25 mM ammonium bicarbonate (ABC) buffer with 1:100 (v/v) EDTA-free protease inhibitor. The cell solution was mixed with 0.1 mm beads and ABC buffer with a ratio of 1:1:2 and lysed by beading for 3 minutes. The cell lysate was centrifuged at 12,000 g (at 4 °C) for 4 min to remove insoluble debris. Subsequently, the supernatant was filtered and concentrated using an Amicon Ultra-0.5 centrifugal filter by centrifuging at 14,000 g (at 4 °C) for 20 min. 1 µL of 1 M dithiothreitol (DTT) was added for reduction at 55 °C for 45 min. The concentration of the lysate was measured using the Pierce bicinchoninic acid (BCA) Protein Assay Kit (Thermo Fisher).

### Top-down RPLC-MS/MS

*E. coli* proteins (300 ng) extracted from the sample were analyzed using an Orbitrap Fusion Lumos mass spectrometer (Thermo Fisher Scientific) coupled with an Ultimate 3000 (Thermo Fisher Scientific) reversed-phase liquid chromatography (RPLC) separation system with a C2 column (100 μm i.d., 60 cm length, CoAnn Inc.). In the RPLC system, phase A was water with 0.1% formic acid (FA), and phase B was 60% acetonitrile (ACN) and 15% isopropanol (IPA) with 0.1% FA. A 98-min gradient of mobile phase B (0-5 min 5%, 5-7 min for 5% to 35%, 7-10 min for 35% to 50%, 10-97 min for 50% to 80%, 97-98 min from 80% to 99%) was applied with a flow rate of 400 nL/min.

*E. coli* proteins were analyzed using both DDA and DIA modes. In each mode, six runs were performed separately, each targeting a specific *m*/*z* range of precursor ions: 720-800, 800-880, 880-960, 960-1040, 1040-1120, and 1120-1200 *m*/*z*. MS1 spectra were collected with the specific *m*/*z* range, a resolution of 240,000 (at 200 *m*/*z*), 4 micro scans, an automatic gain control (AGC) target value of 1x10^6^, and a maximum injection time of 200 ms. MS/MS spectra were obtained with a scan range of 400-2000 *m*/*z*, a resolution of 60,000 (at 200 *m*/*z*), 1 micro scan, an AGC target value of 1×10^6^, and a maximum injection time of 500 ms. Fragmentation was performed using higher-energy collisional dissociation (HCD) with 30% nominal collision energy (NCE). In the DDA runs, the top six precursor ions from each MS1 scan were isolated with a 3 *m*/*z* window for MS/MS analysis, and the dynamic exclusion was set to 60 seconds. In the DIA runs, a 4 *m*/*z* isolation window was used, resulting in a total of 20 MS/MS spectra for each cycle (Supplementary Fig. S1). For the DIA mode, three replicates were obtained: one for method development, and the other two for evaluation. For the DDA mode, two replicates were generated for evaluation. MsConvert^45^ was used to convert raw files into centroided mzML files. Only spectra between 0-113 minutes were kept for further processing.

### Feature extraction for TD-DIA-MS data

MS and MS/MS spectra were preprocessed to remove peaks with intensities below a noise cutoff. We calculated a noise intensity for each MS1 and MS/MS spectrum. To determine the noise intensity cutoff for a spectrum, a histogram of the intensities of all peaks in the spectrum was generated. The noise intensity level *h* was set to the middle value of the bin with the highest frequency^46^. Using a signal/noise (S/N) ratio of *r*_1_ for MS1 spectra and *r*_2_ for MS/MS spectra (*r*_1_ = 3 and *r*_2_ = 1 in the experiments), all peaks with an intensity less than *r*_1_*h* were removed from MS1 spectra, and peaks with an intensity less than *r*_2_*h* were removed from MS/MS spectra.

Proteoform features were extracted from the LC-MS map of each run using a modified version of TopFD^47^ (Supplementary Table S1). A proteoform feature may have isotopic envelopes with different charge states in different scans. The set of all isotopic envelopes of the same charge state of a proteoform feature is called a single charge proteoform feature (SCPF). An SCPF was assigned to an isolation window if the window contained more than 50% of the total peak intensity of the SCPF. As a result, each isolation window has a set of assigned SCPFs.

For each isolation window, an LC-MS/MS map was obtained by combining all MS/MS spectra generated from the window, and a modified version of TopFD was applied to extract fragment features from the LC-MS/MS map (Supplementary Table S2).

### Pseudo spectrum generation

Each TD-DIA-MS run is divided into cycles, each of which contains an MS1 scan and 20 MS/MS scans. All the cycles in a run are sorted in the increasing order of the retention time and the index of a cycle is its position in the sorted list. The extracted ion chromatogram (XIC) of an SCPF or a fragment feature in a TD-DIA-MS run is represented by a vector [*a*_1_, *a*_2_, …, *a*_*k*_], where *a*_*i*_, 1 ≤ *i* ≤ *k*, is the total peak intensity of the SCPF observed in cycle *i*, and *k* is the total number of cycles in the TD-DIA-MS run. The apex cycle distance of an SCPF and a fragment feature is the difference between the cycle indexes of the apex intensity scans of the two features.

For each isolation window, we obtain a list of fragment features extracted from the corresponding LC-MS/MS map and a list of SCPFs assigned to the isolation window. We sort the SCPFs in the list based on their intensities and iteratively generate a pseudo MS/MS spectrum for each SCPF following the order in the sorted list. Once a fragment feature is used in the pseudo MS/MS spectrum of an SCPF, it will be removed from the fragment feature list.

Given an SCPF, three rounds of filtering are performed to shortlist fragment features matched to the SCPF, and the matched fragment features are used to generate a pseudo spectrum. In the first round, a fragment feature is filtered out if the apex cycle distance of the SCPF and the fragment feature > min {*t*, ⌊*c*/2⌋}, where *c* is the number of cycles in which the SCPF is observed, and *t* is a user-specified parameter (*t* =3 in the experiments). The list of remaining fragment features after the first filtering is referred to as *L*.

In the second round, a scoring function is used to filter fragment features. We trained a logistic regression model to assign a score for each fragment feature in *L* using three attributes: a normalized intensity rank of the fragment feature, a normalized cycle number of the fragment feature, and the shared XIC between the SCPF and the fragment feature. The normalized intensity rank is the ratio between the intensity rank in the decreasing order of the fragment feature in *L* and the total number of features in *L*. The normalized cycle number of the fragment feature is the total number of cycles in which the feature is observed divided by the total number of cycles in which the SCPF is observed. To compute the shared XIC, we first perform linear interpolation on the XICs of the SCPF and fragment feature (Supplementary Fig. S2). Subsequently, we normalized the XIC so that the total area under the XIC equals 1. The shared area under the normalized XICs of the SCPF and the fragment feature is reported as the shared XIC. For the SCPF with a set *L* of matched fragment features, we obtain a subset of *L* containing only fragment features with a score greater than a user-specified cutoff (0.55 in the experiments). When the size of the subset < 25, the top 25 fragment features in *L* are reported without filtering.

In the third round, all the remaining fragment features are divided into two groups: a low-mass group (mass < 1500 Da) and a high-mass group (mass ≥ 1500 Da). The top scoring 25 masses in the low mass group and the top scoring *T* - 25 masses in the high mass group are reported, where *T* is the estimated number of possible theoretical N-terminal and C-terminal fragment masses of the precursor ion and 25 is the estimated number of theoretical N-terminal and C-terminal fragment masses with a mass < 1500. To compute *T*, we estimate the proteoform length *l* of the precursor using the precursor mass and the average mass of amino acid residues in the Averagine model^48^ and then set *T* = 2(*l*-1).

### Proteoform Identification

TopFD^47^ was used to analyze the *E. coli* TD-DDA-MS data for proteoform feature detection and spectral deconvolution (Supplementary Table S3), and the pseudo spectrum generation method was employed to produce pseudo MS/MS spectra from the *E. coli* TD-DIA-MS data. The deconvoluted MS/MS spectra or pseudo MS/MS spectra were searched against the UniProt *E. coli* proteome sequence database (4530 entries, version September 06, 2023) using TopPIC^49^ (version 1.7.2). Oxidation, methylation, acetylation, and phosphorylation were selected as variable post-translational modifications (PTMs) (Supplementary Table S4). One unknown mass shift or three variable PTM sites were allowed in one proteoform in the database search. The error tolerances for precursor and fragment masses were set to 10 ppm. Using the target-decoy approach^50^, proteoform identifications were filtered with a 1% FDR cutoff. The parameter settings for TopPIC are provided in Supplementary Table S5.

In postprocessing, duplicated proteoform identifications in each MS run were removed. All proteoform identifications in an MS run were ranked in the decreasing order of the SCPF intensity, and, following the order, each proteoform was compared with the proteoforms with a better rank to check if it was a duplicated one. Two proteoforms with neutral monoisotopic precursor masses *m*_1_ and *m*_2_ identified in an MS run were treated as duplicated ones if the two proteoforms were from the same protein and min{|*m*_1_ − *m*_2_|, |*m*_1_ − *m*_2_ − 1.00235|, |*m*_1_ − *m*_2_ + 1.00235|} is no more than 10 ppm, where 1.00235 Da is a common error in top-down spectral deconvolution. For each identified duplicated proteoform pair, the proteoform with a lower SCPF intensity was removed from the proteoform list. To combine proteoforms identified from the six MS/MS runs with different *m*/*z* ranges for precursor ions for a protein sample, we first merged the identified proteoforms in the six runs and then used the above method to remove duplicated proteoforms.

## Results

### Overview of the TopDIA pipeline

The TopDIA pipeline is shown in Fig. 1. In the pipeline, we first identify proteoform features and SCPFs in the LC-MS map. For each isolation window, we assign a list of SCPFs to the window based on the overlap of the isolation window and isotopic peaks of the SCPF (Methods) and extract fragment features in the corresponding LC-MS/MS map. Then we rank the SCPFs assigned to the window in the decreasing order of the total peak intensity. For the best-ranking SCPF, all fragment features of the isolation window are filtered using a three-step method (Methods) and the remaining fragment features and the SCPF are used to generate a pseudo MS/MS spectrum. Then the matched fragment features are removed from the fragment feature list and then the second best SCPF is selected to generate a pseudo MS/MS spectrum. The pseudo spectra generation step is repeated for all the SCPFs assigned to the window following the decreasing order of the total peak intensity. The resulting pseudo spectra are searched against a protein sequence database for proteoform identification using TopPIC^49^ (Methods).

**Figure 1.**
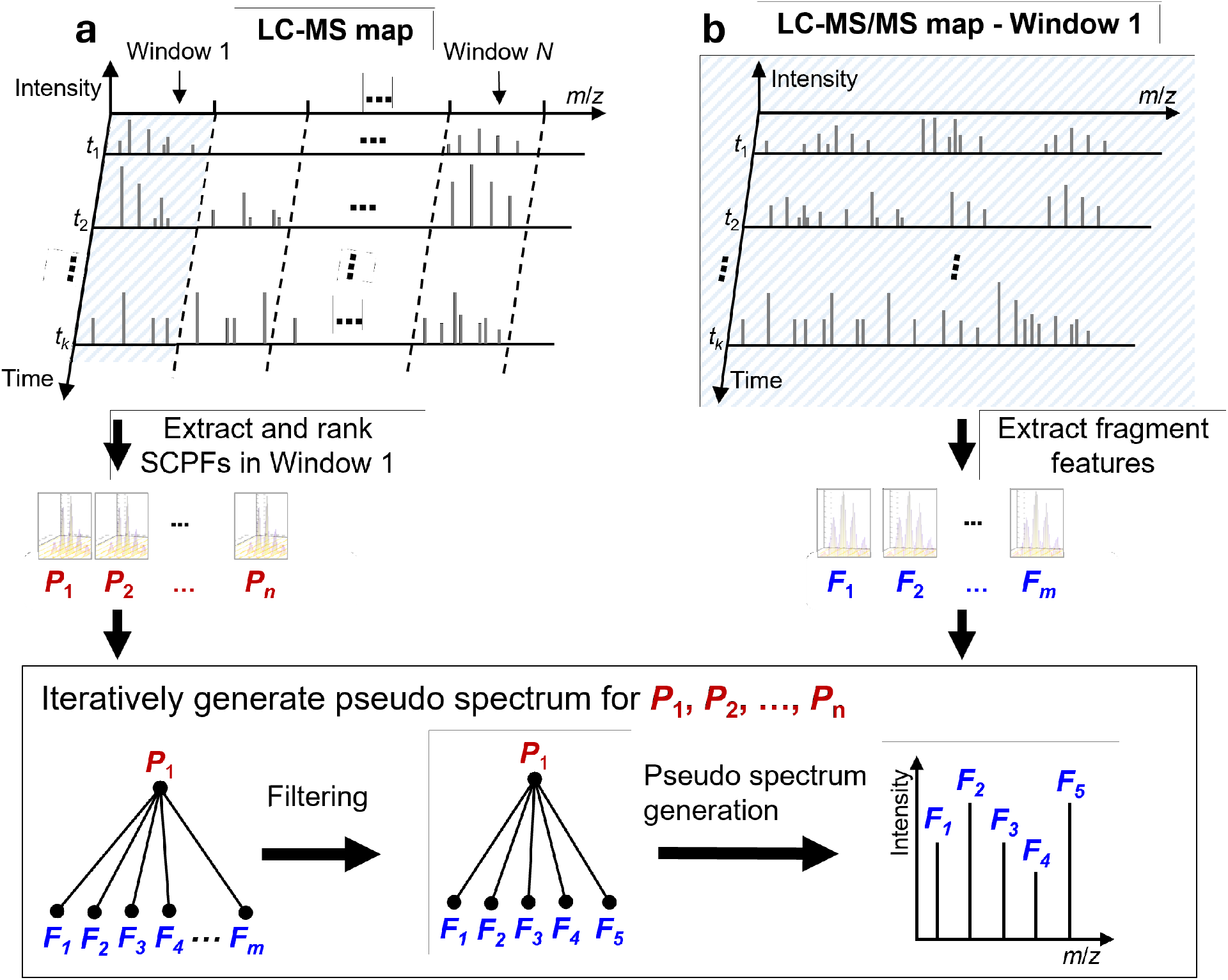
Generation of pseudo MS/MS spectra using an isolation window (window 1) in an LC TD-DIA-MS data. (a) Proteoform features detected in the LC-MS map and SCPFs are assigned to isolation window 1 and then ranked in the decreasing order of the total peak intensity. (b) Fragment features are extracted from the LC-MS/MS map of the isolation window 1. Pseudo spectra were iteratively generated with the order *P*_1_, *P*_2_, …, *P*_n_. For SCPF *P*_1_, all fragment features *F*_1_, …, *F*_m_, are filtered, and the SCPF *P*_1_ and the remaining fragment features are used to construct a pseudo MS/MS spectrum for *P*_1_. Then the fragment features used in the pseudo spectrum are removed from the fragment feature list, and the second SCPF *P*_2_ is selected to generate a pseudo spectrum using the same method.

### A logistic regression model for filtering fragment features

A logistic regression model was trained to evaluate the match between an SCPF and a fragment feature. As TD-DIA-MS was slow to cover a large *m*/*z* range with small isolation windows in a cycle, the *m*/*z* range [720, 1200] was divided into six 80 *m*/*z* ranges. Three replicates of TD-DIA-MS data were generated from *E. coli* K-12 MG1655 cells, and each replicate contained 6 MS runs (Methods). The first replicate, referred to as DIA-TRAIN, was used to train the logistics regression model. The DIA-TRAIN data was analyzed using the TopPIC suite pipeline^51^, in which TopFD was used for spectral deconvolution and proteoform feature detection (see Supplementary Table S3 for parameter settings of TopFD) and TopPIC was employed to search the deconvoluted MS/MS spectra against the UniProt *E. coli* proteome sequence database for proteoform identification (see Supplementary Table S5 for parameter settings of TopPIC). Only identified proteoforms without variable modifications and unknown modifications (434 proteoforms) were kept.

We found all SCPFs of the 434 proteoforms and assigned each SCPF to an isolation window (See Methods). Additionally, we acquired fragment features for each isolation window of each run using a modified version of TopFD (Supplementary Table S2). A fragment feature was paired to an SCPF if they were from the same isolation window and their apex cycle distance was less than a cutoff value (see Methods). Note that a fragment feature could be paired with multiple SCPFs. The fragment feature and SCPF pairs were labeled positive if the fragment feature matched a b- or y-ion mass of the proteoform identification, and negative otherwise. In total, we identified 75,852 SCPF and fragment feature pairs, of which 10,280 were labeled positive and 65,572 were negative. We randomly split the labeled data with a 70:30 ratio into training and validation sets. We trained the model using the training set and achieved a balanced accuracy of 78.14% and the area under the receiver operating characteristic (ROC) curve (AUC) value of 84.78% on the validation data set (Supplementary Fig. S3).

## Comparison of DDA and DIA

Three replicates of TD-DIA-MS data and two replicates of TD-DDA-MS data were generated from *E. coli* K-12 MG1655 cells (see Methods). The second and third replicates of the TD-DIA-MS data referred to as DIA-TEST-1 and DIA-TEST-2, and the two replicates of the TD-DDA-MS data, referred to as DDA-TEST-1 and DDA-TEST-2, were used to compare proteoform identifications of the two approaches. The TopPIC suite pipeline^51^ and the proposed TopDIA pipeline were used to search the DDA and DIA MS data against the UniProt *E. coli* proteome database for proteoform identification, respectively (see Methods).

With a 1% proteoform level FDR cutoff, TD-DIA-MS increased the average number of proteoform identifications reported in two replicates from 574 to 627.5 and increased the average number of protein identifications from 186 to 205.5 compared with TD-DDA-MS. However, for most single MS runs, TD-DDA-MS reported more proteoforms and proteins than TD-DIA-MS (Fig. 2a and 2b, and Supplementary Tables S6 and S7). The reason was that proteoform identifications reported by different *m*/*z* ranges of the TD-DIA-MS runs had fewer overlapping identifications compared with the TD-DDA-MS runs.

**Figure 2.**
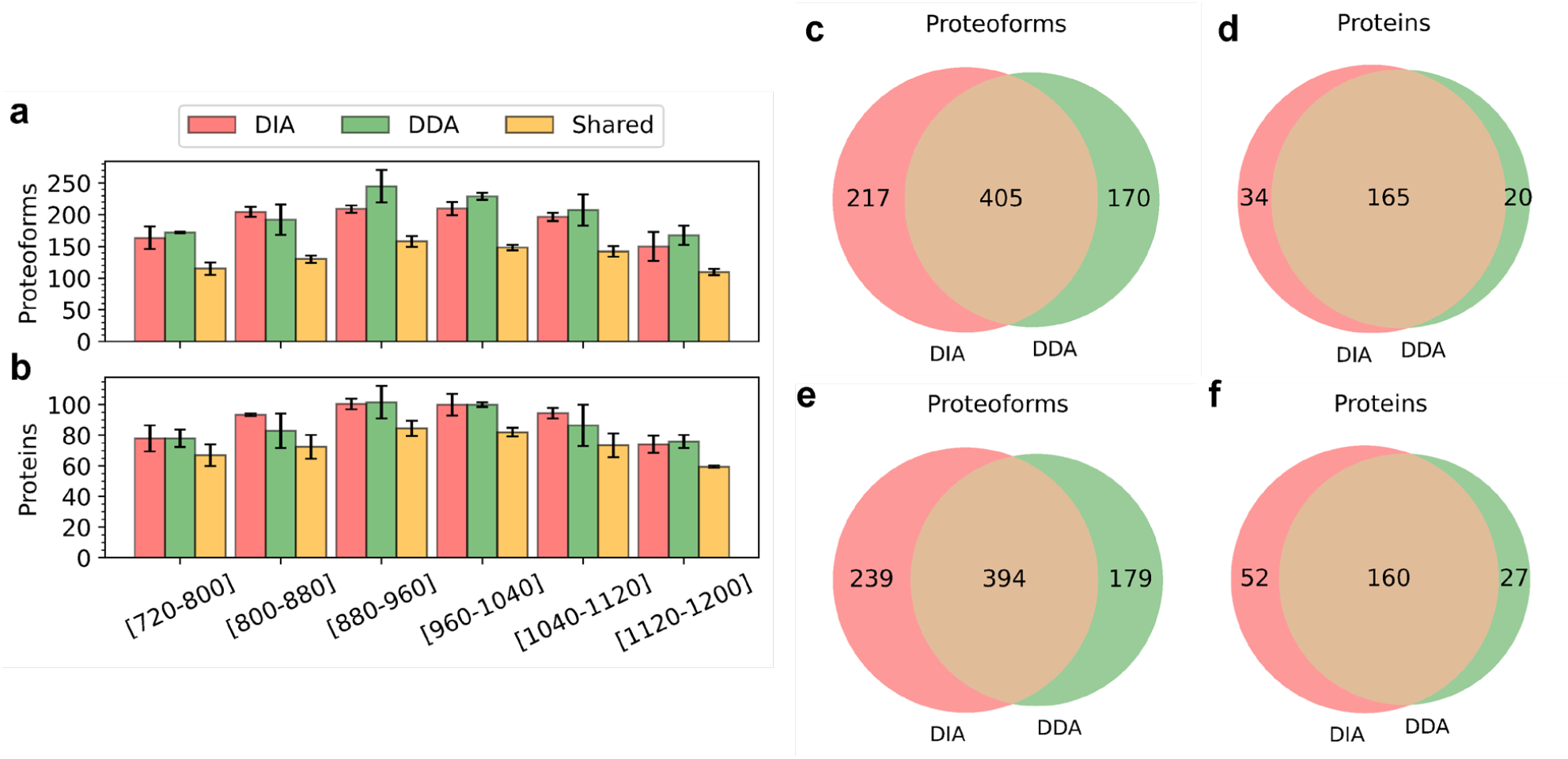
Comparison of TD-DIA-MS and TD-DDA-MS on proteoform and protein identification. Comparison of the average numbers of proteoforms (a) and proteins (b) identified from the six runs of DIA-TEST-1 and DIA-TEST-2 and those from DDA-TEST-1 and DDA-TEST-2. Venn diagrams showing the overlaps of proteoforms (c), and proteins (d) identified from DIA-TEST-1 and DDA-TEST-1 as well as the overlaps of proteoforms (e), and proteins (f) identified from DIA-TEST-2 and DDA-TEST-2.

The average number of proteoform and protein identifications shared by the two approaches were 399.5 and 162.5, respectively (Fig. 2c-2f). There were three main reasons why TD-DDA-MS missed some proteoform identifications reported by TD-DIA-MS. First, some low-intensity proteoform features were not selected for MS/MS analysis in TD-DDA-MS. Second, some DDA MS/MS spectra were multiplexed, and the TopPIC pipeline often failed to identify any proteoforms from these multiplexed spectra. Even if TopPIC identified the proteoform for the most intense SCPF, it missed proteoform identifications of other SCPFs of the multiplexed spectrum. Third, some DDA MS/MS spectra lacked sufficient fragment ions for confident proteoform identification. TD-DIA-MS also missed some proteoform identifications reported by TD-DDA-MS due to the low frequency for MS1 spectral acquisition and limitations in the identification of multiplexed MS/MS spectra. A total of 20 MS/MS scans were acquired in each TD-DIA-MS cycle, which needed more time for spectral acquisition than 6 MS/MS scans in each TD-DDA-MS cycle. As a result, MS1 scans in the TD-DIA-MS data failed to provide high-quality features for some proteoforms, leading to missed identifications. In addition, TopDIA sometimes failed to detect and assign enough fragment masses to some SCPFs, resulting in missing identifications. For example, some multiplexed MS/MS spectra were dominated by fragment ions from one SCPF, making it impossible to detect enough fragment masses for other low-intensity SCPFs.

We compared the fragment masses in pseudo MS/MS spectra reported by TopDIA from DIA-TEST-1 and those in single MS/MS spectra reported by TopFD from DDA-TEST-1. A total of 405 proteoforms were identified by both data sets. For each of the 405 proteoforms, we found the pseudo spectrum with the best PrSM *E*-value from DIA-TEST-1 and the single MS/MS spectrum with the best PrSM *E*-value from DDA-TEST-1. The selected DIA spectra had an average of 45.66 fragment mass per spectrum, which was less than that (91.01) of the selected DDA spectra. But the average number of matched fragment masses for the DIA pseudo spectra was only slightly less than that of the DDA MS/MS spectra (DIA: 17.76 vs DDA: 20.89), showing that the DIA spectra contained a higher percentage of b- and y-fragment masses than the DDA spectra (38.89% vs 22.96%). A similar pattern was observed in the DIA-TEST-2 and DDA-TEST-2 data sets (Fig. 3).

**Figure 3.**
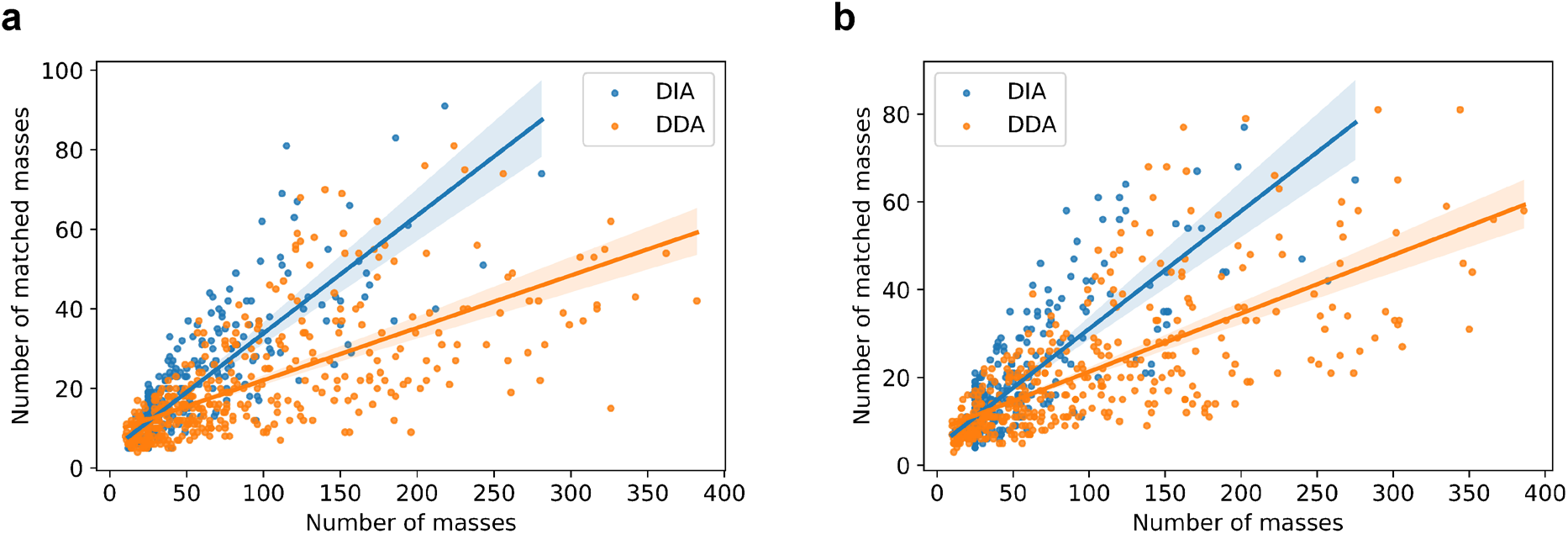
The numbers of fragment masses and matched fragment masses in the best DIA pseudo spectrum and the best DDA single spectrum for each of the proteoform identifications shared by the *E. coli* TD-DIA-MS and TD-DDA-MS data. (a) DIA-TEST-1 vs DDA-TEST-2, and (b) DIA-TEST-2 vs DDA-TEST-2.

### Comparison of pseudo spectra and single spectra in TD-DIA-MS data

We compared pseudo spectra reported by TopDIA and single MS/MS spectra on proteoform identification using the DIA-TEST-1 data set. As many single MS/MS spectra in the data are dominated by fragment ions of one SCPF, these spectra can be treated as non-multiplexed ones for proteoform identification. In the single spectra approach, TopFD was utilized for spectral deconvolution and proteoform feature detection (Supplementary Table S3), and TopPIC was employed to search the deconvoluted MS/MS spectra against the UniProt E. coli proteome sequence database to identify proteoforms (Supplementary Table S5). In the pseudo spectra approach, TopDIA was used to generate pseudo-MS/MS spectra, followed by proteoform identification using TopPIC (see Methods). In all six runs, the pseudo spectra approach reported more protein and proteoform identifications than the single spectra approach (Fig. 4a and 4b, and Supplementary Table S8). After merging the identifications from the six runs, the pseudo spectra approach increased proteoform identifications by 12.27% (622 vs 554) and protein identifications by 8.15% (199 vs 184) compared with the single spectra approach (Fig. 4c and 4d). The pseudo spectra approach also missed some proteoforms reported by the single spectra approach. The reason was that many fragment masses were filtered out during the generation of pseudo spectra for some SCPFs. Consequently, the resulting pseudo spectra lacked sufficient fragment masses for proteoform identification.

**Figure 4.**
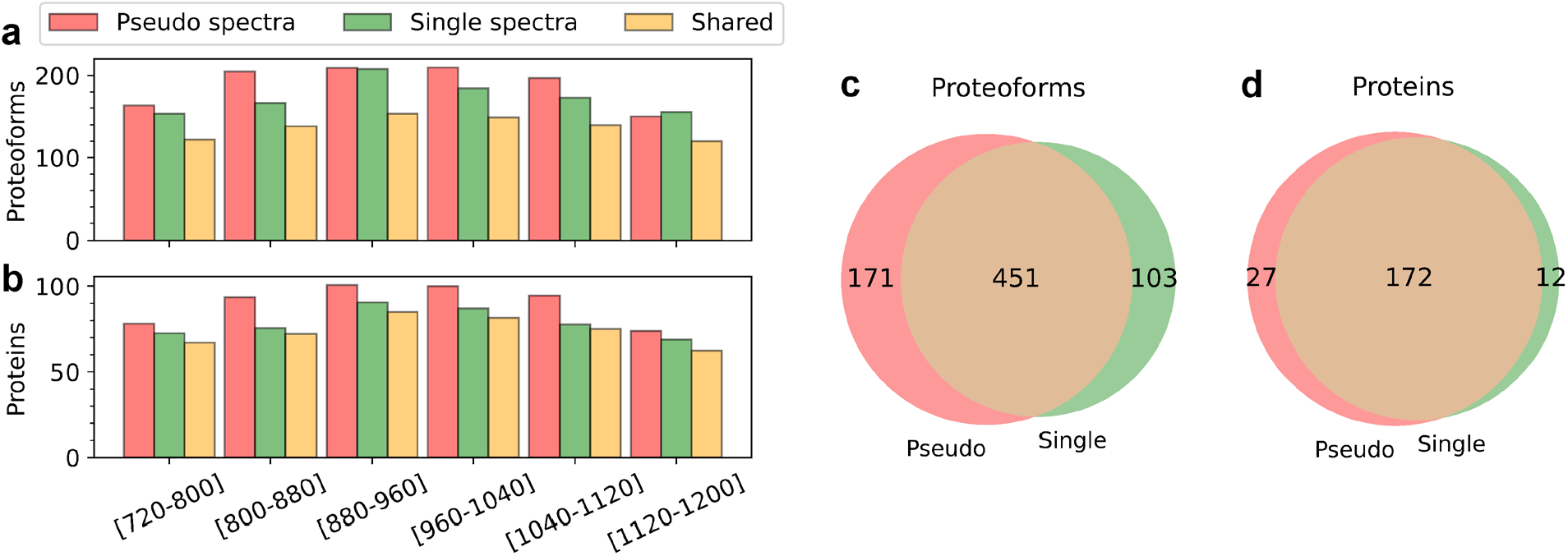
Comparison of the pseudo spectra approach and single spectra approach for proteoform and protein identification in TD-DIA-MS. Comparison of the number of proteoforms (a), and proteins (b) identified from the six runs of DIA-TEST-1 using the two approaches. Venn diagrams showing the overlaps of the total proteoforms (c), and proteins (d) identified from the six runs of DIA-TEST-1 using the two approaches.

### Reproducibility across technical replicates

We compared the reproducibility of proteoform identifications reported from DIA-TEST-1 and DIA-TEST-2 with that of DDA-TEST-1 and DDA-TEST-2. For an *m*/*z* range, e.g. [720-800], let *n*_1_ and *n*_2_ be the numbers of proteoforms reported from DIA-TEST-1 and DDA-TEST-1, respectively. We sorted the proteoform identifications in the increasing order of *E*-values and then kept only the best *n* = min {*n*_1_, *n*_2_} proteoforms reported from each data set. The same method was applied to filter proteoform identifications reported from the runs with the same *m*/*z* range in DIA-TEST-2 and DDA-TEST-2. Then the overlap coefficient of proteoform identifications of the two DIA replicates was compared with that of the two DDA replicates. The same method was applied to compare the overlaps of the proteins and proteoforms reported for each *m*/*z* range from the two DIA replicates and two DDA replicates (Fig. 5a and 5b, and Supplementary Tables S9 and S10). Using the method, we found that TD-DDA-MS achieved higher overlap coefficients for protein and proteoform identifications than TD-DIA-MS for all six *m*/z ranges. The lower overlap coefficients of TD-DIA-MS can be attributed to its lower frequency in MS1 spectral acquisition.

**Figure 5.**
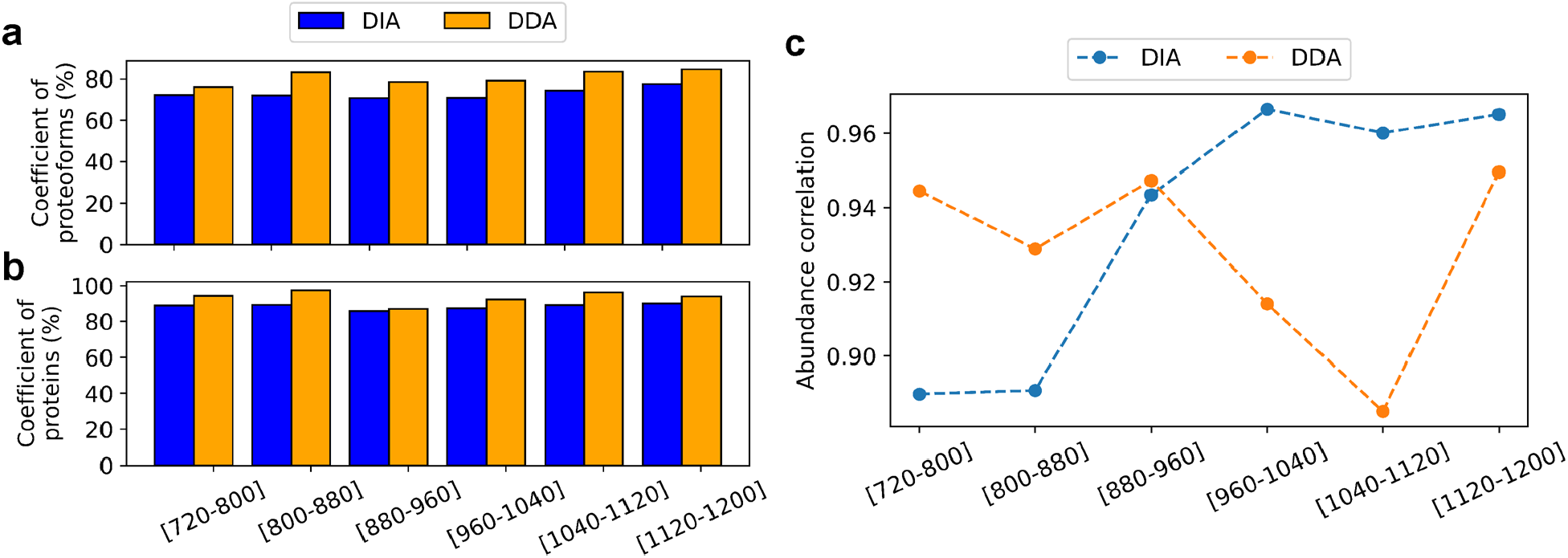
Comparison of the overlap coefficients of proteoforms (a), and proteins (b) identified in DIA-TEST-1 and DIA-TEST-2 with those of DDA-TEST-1 and DDA-TEST-2. (c) Quantitative reproducibility of common proteoform identified in the four data sets.

After merging the identifications from the six runs, we filtered the proteoforms to report the same number of identifications from DIA-TEST-1 and DDA-TEST-1 and the same number of identifications from DIA-TEST-2 and DDA-TEST-2. TD-DDA-MS reported a higher overlap coefficient compared to TD-DIA-MS at the proteoform level (DIA: 70.51% vs DDA: 74.69%) and protein level (DIA: 85.48% vs DDA: 88.65%). We also examined the reproducibility without proteoform filtering. TD-DIA-MS reported 432 proteoforms with an overlap efficiency of 69.45%, and TD-DDA-MS reported 428 proteoforms with an overlap efficiency of 74.69% from two replicates without filtering. TD-DIA-MS and TD-DDA-MS identified the same number of proteins (164) without filtering with overlapping coefficients of 82.41% and 88.64%, respectively.

We also compared TD-DIA-MS and TD-DDA-MS in the reproducibility of proteoform quantification. For each of the six *m*/*z* ranges, we selected the proteoforms identified from all four data sets, and then computed the Pearson correlation coefficient (PCC) of the logarithm (base 2) transformed abundances of the selected proteoforms for the DIA and DDA runs (Fig. 5c). The DIA data reported better PCCs in three runs and worse PCCs in the other three runs compared with the DDA data. After merging the identifications from the six runs, we obtained 314 proteoforms identified from all four data sets. TD-DDA-MS reported a PCC value of 0.9056 for logarithm-transformed proteoform abundances while TD-DIA-MS obtained a similar PCC value of 0.9023.

## Discussion

There are still many challenges in proteoform identification by TD-DIA-MS. The main challenge is the low spectral acquisition frequency. Top-down MS usually relies on Orbitrap or Fourier-transform ion cyclotron resonance (FT-ICR) mass spectrometers for spectral acquisition, whose spectral acquisition frequencies are low for collecting high-resolution spectra. For example, the scan speed is about 7.5 Hz for a Thermo Orbitrap with a spectral resolution of 60K, and several seconds are needed to collect 20 MS/MS spectra in each DIA cycle. As a result, only one or two MS1 scans are collected for proteoforms with a short elution profile, and the MS1 scan may fail to provide high-quality proteoform features, leading to missing identifications of the proteoforms.

Another challenge is the complexity of top-down multiplexed MS/MS spectra, which contain many more fragment ion peaks than bottom-up multiplexed MS/MS spectra. Because of this, isotopic envelopes of fragment ions often overlap with each other in top-down multiplexed MS/MS spectra and low-abundance fragment ions may be not detected, resulting in missing identifications of proteoforms.

With gas phase fractionation, a small *m*/*z* range of precursor ions can be selected to reduce the MS/MS scan numbers in each DIA cycle. In the TD-DIA-MS analysis of *E. coli* proteins, each MS run covered an 80 *m*/*z* range, and a small 4 *m*/*z* isolation window was selected to reduce the complexity of DIA MS/MS spectra. Using this MS experiment setting, TD-DIA-MS achieved similar performance in proteoform identification and quantification compared with TD-DDA-MS. However, a total of 6 MS runs were used to cover the *m*/*z* range [720, 1200], making TD-DIA-MS experiments time-consuming.

Characterization of proteoforms with PTMs or unknown mass shifts is also a challenging problem in TD-DIA-MS. For multiplexed DIA MS/MS spectra containing fragment ions from two proteoforms of two different proteins, fragment ions of the second proteoform may introduce errors in the characterization of the first proteoform. When a multiplexed MS/MS spectrum is generated from two proteoforms of the same protein, many fragment ions in the spectrum are shared by the two proteoforms, and proteoform characterization relies on only fragment ions that are unique for each proteoform^52^.

The reproducibility in proteoform identification and quantification by TD-DIA-MS is limited by its low spectral acquisition frequency. The experimental results showed that the reproducibility of TD-DIA-MS was slightly worse than TD-DDA-MS in proteoform identification and quantification. Increasing the speed of high-resolution spectral acquisition is essential to further improvement of the reproducibility of TD-DIA-MS.

The isolation window size and gas phase fractionation are important experiment settings in TD-DIA-MS. Experimental results showed that small isolation windows are needed to reduce the complexity of top-down DIA MS/MS spectra. Due to the speed limitation in spectral acquisition, gas phase fractionation is required to cover the *m*/*z* range of MS1 scans using small isolation windows.

## Conclusions

In this paper, we presented TopDIA, the first software tool for proteoform identification by demultiplexing top-down DIA MS/MS spectra. TopDIA generates pseudo non-multiplexed MS/MS spectra from TD-DIA-MS data by integrating algorithms for detecting and matching proteoform and fragment features. Experimental results on the *E. coli* data demonstrated that TD-DIA-MS with TopDIA increased proteoform and protein identifications compared with TD-DDA-MS when gas phase fractionation with multiple MS runs was employed. We further demonstrated that pseudo spectra reported by TopDIA from TD-DIA-MS data contained a higher percentage of fragment masses matched to identified proteoforms compared with single MS/MS spectra in TD-DDA-MS data. Assigning fragment features to precursor features is the main challenging computational problem in proteoform identification by TD-DIA-MS. A future research direction is to employ deep-learning models to further improve the accuracy in the generation of pseudo spectra, which could enhance proteoform identifications by TD-DIA-MS.

## Data Availability

TopDIA is available as part of the TopPIC suite at https://github.com/toppic-suite/toppic-suite/releases/tag/v1.7_DIA. The *E. coli* data set is available at the MassIVE repository (ID: MSV000094407). The data and Python scripts for training the logistic regression model and evaluating the identification performance of TopDIA are available at https://www.toppic.org/software/toppic/topdia_supplemental.html.

## Acknowledgments

This research was funded by NIH through the grants R01GM118470 and R01CA247863.

## Notes

### Competing Interest Statement

The authors have declared no competing interest.

## References

(1) Donnelly, D. P.; Rawlins, C. M.; DeHart, C. J.; Fornelli, L.; Schachner, L. F.; Lin, Z.; Lippens, J. L.; Aluri, K. C.; Sarin, R.; Chen, B. Best practices and benchmarks for intact protein analysis for top-down mass spectrometry. Nature Methods 2019, 16 (7), 587–594. DOI: 10.1038/s41592-019-0457-0.

(2) Cui, W.; Rohrs, H. W.; Gross, M. L. Top-down mass spectrometry: recent developments, applications and perspectives. Analyst 2011, 136 (19), 3854–3864. DOI: 10.1039/C1AN15286F.

(3) Brown, K. A.; Melby, J. A.; Roberts, D. S.; Ge, Y. Top-down proteomics: challenges, innovations, and applications in basic and clinical research. Expert review of proteomics 2020, 17 (10), 719–733. DOI: 10.1080/14789450.2020.1855982.

(4) McCool, E. N.; Xu, T.; Chen, W.; Beller, N. C.; Nolan, S. M.; Hummon, A. B.; Liu, X.; Sun, L. Deep top-down proteomics revealed significant proteoform-level differences between metastatic and nonmetastatic colorectal cancer cells. Science Advances 2022, 8 (51), eabq6348. DOI: 10.1126/sciadv.abq6348.

(5) Melani, R. D.; Gerbasi, V. R.; Anderson, L. C.; Sikora, J. W.; Toby, T. K.; Hutton, J. E.; Butcher, D. S.; Negrão, F.; Seckler, H. S.; Srzentić, K. The Blood Proteoform Atlas: A reference map of proteoforms in human hematopoietic cells. Science 2022, 375 (6579), 411–418. DOI: 10.1126/science.aaz5284.

(6) Tabb, D. L.; Vega-Montoto, L.; Rudnick, P. A.; Variyath, A. M.; Ham, A.-J. L.; Bunk, D. M.; Kilpatrick, L. E.; Billheimer, D. D.; Blackman, R. K.; Cardasis, H. L.; et al. Repeatability and Reproducibility in Proteomic Identifications by Liquid Chromatography™Tandem Mass Spectrometry. Journal of Proteome Research 2010, 9 (2), 761–776. DOI: 10.1021/pr9006365.

(7) Bantscheff, M.; Lemeer, S.; Savitski, M. M.; Kuster, B. Quantitative mass spectrometry in proteomics: critical review update from 2007 to the present. Analytical and bioanalytical chemistry 2012, 404 (4), 939–965. DOI: 10.1007/s00216-012-6203-4.

(8) Yergey, A. L.; Edmonds, C. G.; Lewis, I. A.; Vestal, M. L. Liquid chromatography/mass spectrometry: techniques and applications; Springer Science & Business Media, 2013. DOI: 10.1007/978-1-4899-3605-9.

(9) Makarov, A.; Scigelova, M. Coupling liquid chromatography to Orbitrap mass spectrometry. Journal of Chromatography A 2010, 1217 (25), 3938–3945. DOI: 10.1016/j.chroma.2010.02.022.

(10) Chen, D.; McCool, E. N.; Yang, Z.; Shen, X.; Lubeckyj, R. A.; Xu, T.; Wang, Q.; Sun, L. Recent advances (2019–2021) of capillary electrophoresis-mass spectrometry for multilevel proteomics. Mass Spectrometry Reviews 2023, 42 (2), 617–642. DOI: 10.1002/mas.21714.

(11) Lubeckyj, R. A.; Basharat, A. R.; Shen, X.; Liu, X.; Sun, L. Large-Scale Qualitative and Quantitative Top-Down Proteomics Using Capillary Zone Electrophoresis-Electrospray Ionization-Tandem Mass Spectrometry with Nanograms of Proteome Samples. Journal of the American Society for Mass Spectrometry 2019, 30 (8), 1435–1445. DOI: 10.1007/s13361-019-02167-w.

(12) Nesvizhskii, A. I. A survey of computational methods and error rate estimation procedures for peptide and protein identification in shotgun proteomics. Journal of Proteomics 2010, 73 (11), 2092–2123. DOI: 10.1016/j.jprot.2010.08.009.

(13) Bilbao, A.; Varesio, E.; Luban, J.; Strambio-De-Castillia, C.; Hopfgartner, G.; Müller, M.; Lisacek, F. Processing strategies and software solutions for data-independent acquisition in mass spectrometry. PROTEOMICS 2015, 15 (5-6), 964–980. DOI: 10.1002/pmic.201400323.

(14) Bruderer, R.; Bernhardt, O. M.; Gandhi, T.; Miladinović, S. M.; Cheng, L.-Y.; Messner, S.; Ehrenberger, T.; Zanotelli, V.; Butscheid, Y.; Escher, C.; et al. Extending the Limits of Quantitative Proteome Profiling with Data-Independent Acquisition and Application to Acetaminophen-Treated Three-Dimensional Liver Microtissues. Molecular & Cellular Proteomics 2015, 14 (5), 1400–1410. DOI: 10.1074/mcp.M114.044305.

(15) Bailey, D. J.; McDevitt, M. T.; Westphall, M. S.; Pagliarini, D. J.; Coon, J. J. Intelligent Data Acquisition Blends Targeted and Discovery Methods. Journal of Proteome Research 2014, 13 (4), 2152–2161. DOI: 10.1021/pr401278j.

(16) Michalski, A.; Cox, J.; Mann, M. More than 100,000 Detectable Peptide Species Elute in Single Shotgun Proteomics Runs but the Majority is Inaccessible to Data-Dependent LC™MS/MS. Journal of Proteome Research 2011, 10 (4), 1785–1793. DOI: 10.1021/pr101060v.

(17) Ludwig, C.; Gillet, L.; Rosenberger, G.; Amon, S.; Collins, B. C.; Aebersold, R. Data-independent acquisition-based SWATH-MS for quantitative proteomics: a tutorial. Molecular Systems Biology 2018, 14 (8), e8126. DOI: 10.15252/msb.20178126.

(18) Venable, J. D.; Dong, M.-Q.; Wohlschlegel, J.; Dillin, A.; Yates III, J. R. Automated approach for quantitative analysis of complex peptide mixtures from tandem mass spectra. Nature Methods 2004, 1 (1), 39–45. DOI: 10.1038/nmeth705.

(19) Gillet, L. C.; Navarro, P.; Tate, S.; Röst, H.; Selevsek, N.; Reiter, L.; Bonner, R.; Aebersold, R. Targeted Data Extraction of the MS/MS Spectra Generated by Data-independent Acquisition: A New Concept for Consistent and Accurate Proteome Analysis. Molecular & Cellular Proteomics 2012, 11 (6). DOI: 10.1074/mcp.O111.016717.

(20) Zhang, Y.; Fonslow, B. R.; Shan, B.; Baek, M.-C.; Yates III, J. R. Protein analysis by shotgun/bottom-up proteomics. Chemical Reviews 2013, 113 (4), 2343–2394. DOI: 10.1021/cr3003533.

(21) Lou, R.; Cao, Y.; Li, S.; Lang, X.; Li, Y.; Zhang, Y.; Shui, W. Benchmarking commonly used software suites and analysis workflows for DIA proteomics and phosphoproteomics. Nature Communications 2023, 14 (1), 94. DOI: 10.1038/s41467-022-35740-1.

(22) Zhang, F.; Ge, W.; Ruan, G.; Cai, X.; Guo, T. Data-Independent Acquisition Mass Spectrometry-Based Proteomics and Software Tools: A Glimpse in 2020. PROTEOMICS 2020, 20 (17-18), 1900276. DOI: 10.1002/pmic.201900276.

(23) Röst, H. L.; Rosenberger, G.; Navarro, P.; Gillet, L.; Miladinović, S. M.; Schubert, O. T.; Wolski, W.; Collins, B. C.; Malmström, J.; Malmström, L.; et al. OpenSWATH enables automated, targeted analysis of data-independent acquisition MS data. Nature Biotechnology 2014, 32 (3), 219–223. DOI: 10.1038/nbt.2841.

(24) MacLean, B.; Tomazela, D. M.; Shulman, N.; Chambers, M.; Finney, G. L.; Frewen, B.; Kern, R.; Tabb, D. L.; Liebler, D. C.; MacCoss, M. J. Skyline: an open source document editor for creating and analyzing targeted proteomics experiments. Bioinformatics 2010, 26 (7), 966–968. DOI: 10.1093/bioinformatics/btq054.

(25) Peckner, R.; Myers, S. A.; Jacome, A. S. V.; Egertson, J. D.; Abelin, J. G.; MacCoss, M. J.; Carr, S. A.; Jaffe, J. D. Specter: linear deconvolution for targeted analysis of data-independent acquisition mass spectrometry proteomics. Nat Methods 2018, 15 (5), 371–378. DOI: 10.1038/nmeth.4643.

(26) Searle, B. C.; Lawrence, R. T.; MacCoss, M. J.; Villén, J. Thesaurus: quantifying phosphopeptide positional isomers. Nat Methods 2019, 16 (8), 703–706. DOI: 10.1038/s41592-019-0498-4.

(27) Schubert, O. T.; Gillet, L. C.; Collins, B. C.; Navarro, P.; Rosenberger, G.; Wolski, W. E.; Lam, H.; Amodei, D.; Mallick, P.; MacLean, B.; et al. Building high-quality assay libraries for targeted analysis of SWATH MS data. Nature Protocols 2015, 10 (3), 426–441. DOI: 10.1038/nprot.2015.015.

(28) Meyer, J. G.; Mukkamalla, S.; Steen, H.; Nesvizhskii, A. I.; Gibson, B. W.; Schilling, B. PIQED: automated identification and quantification of protein modifications from DIA-MS data. Nature Methods 2017, 14 (7), 646–647. DOI: 10.1038/nmeth.4334.

(29) Tiwary, S.; Levy, R.; Gutenbrunner, P.; Salinas Soto, F.; Palaniappan, K. K.; Deming, L.; Berndl, M.; Brant, A.; Cimermancic, P.; Cox, J. High-quality MS/MS spectrum prediction for data-dependent and data-independent acquisition data analysis. Nature Methods 2019, 16 (6), 519–525. DOI: 10.1038/s41592-019-0427-6.

(30) Zhou, X.-X.; Zeng, W.-F.; Chi, H.; Luo, C.; Liu, C.; Zhan, J.; He, S.-M.; Zhang, Z. pDeep: Predicting MS/MS Spectra of Peptides with Deep Learning. Analytical Chemistry 2017, 89 (23), 12690–12697. DOI: 10.1021/acs.analchem.7b02566.

(31) Gessulat, S.; Schmidt, T.; Zolg, D. P.; Samaras, P.; Schnatbaum, K.; Zerweck, J.; Knaute, T.; Rechenberger, J.; Delanghe, B.; Huhmer, A.; et al. Prosit: proteome-wide prediction of peptide tandem mass spectra by deep learning. Nature Methods 2019, 16 (6), 509–518. DOI: 10.1038/s41592-019-0426-7.

(32) Yang, Y.; Liu, X.; Shen, C.; Lin, Y.; Yang, P.; Qiao, L. In silico spectral libraries by deep learning facilitate data-independent acquisition proteomics. Nature Communications 2020, 11 (1), 146. DOI: 10.1038/s41467-019-13866-z.

(33) Tsou, C.-C.; Avtonomov, D.; Larsen, B.; Tucholska, M.; Choi, H.; Gingras, A.-C.; Nesvizhskii, A. I. DIA-Umpire: comprehensive computational framework for data-independent acquisition proteomics. Nature Methods 2015, 12 (3), 258–264. DOI: 10.1038/nmeth.3255.

(34) Li, Y.; Zhong, C.-Q.; Xu, X.; Cai, S.; Wu, X.; Zhang, Y.; Chen, J.; Shi, J.; Lin, S.; Han, J. Group-DIA: analyzing multiple data-independent acquisition mass spectrometry data files. Nature Methods 2015, 12 (12), 1105–1106. DOI: 10.1038/nmeth.3593.

(35) He, Q.; Zhong, C.-Q.; Li, X.; Guo, H.; Li, Y.; Gao, M.; Yu, R.; Liu, X.; Zhang, F.; Guo, D.; et al. Dear-DIA XMBD: Deep Autoencoder Enables Deconvolution of Data-Independent Acquisition Proteomics. Research 2023, 6, 0179. DOI: 10.34133/research.0179.

(36) Ting, Y. S.; Egertson, J. D.; Bollinger, J. G.; Searle, B. C.; Payne, S. H.; Noble, W. S.; MacCoss, M. J. PECAN: library-free peptide detection for data-independent acquisition tandem mass spectrometry data. Nature Methods 2017, 14 (9), 903–908. DOI: 10.1038/nmeth.4390.

(37) Ting, Y. S.; Egertson, J. D.; Payne, S. H.; Kim, S.; MacLean, B.; Käll, L.; Aebersold, R.; Smith, R. D.; Noble, W. S.; MacCoss, M. J. Peptide-centric proteome analysis: an alternative strategy for the analysis of tandem mass spectrometry data. Molecular & Cellular Proteomics 2015, 14 (9), 2301–2307. DOI: 10.1074/mcp.O114.047035.

(38) Navarro, P.; Kuharev, J.; Gillet, L. C.; Bernhardt, O. M.; MacLean, B.; Röst, H. L.; Tate, S. A.; Tsou, C.-C.; Reiter, L.; Distler, U.; et al. A multicenter study benchmarks software tools for label-free proteome quantification. Nature Biotechnology 2016, 34 (11), 1130–1136. DOI: 10.1038/nbt.3685.

(39) Demichev, V.; Messner, C. B.; Vernardis, S. I.; Lilley, K. S.; Ralser, M. DIA-NN: neural networks and interference correction enable deep proteome coverage in high throughput. Nature Methods 2020, 17 (1), 41–44. DOI: 10.1038/s41592-019-0638-x.

(40) Sinitcyn, P.; Hamzeiy, H.; Salinas Soto, F.; Itzhak, D.; McCarthy, F.; Wichmann, C.; Steger, M.; Ohmayer, U.; Distler, U.; Kaspar-Schoenefeld, S.; et al. MaxDIA enables library-based and library-free data-independent acquisition proteomics. Nature Biotechnology 2021, 39 (12), 1563–1573. DOI: 10.1038/s41587-021-00968-7.

(41) Smith, L. M.; Kelleher, N. L. Proteoform: a single term describing protein complexity. Nature Methods 2013, 10 (3), 186–187. DOI: 10.1038/nmeth.2369.

(42) Catherman, A. D.; Skinner, O. S.; Kelleher, N. L. Top down proteomics: facts and perspectives. Biochemical and biophysical research communications 2014, 445 (4), 683–693. DOI: 10.1016/j.bbrc.2014.02.041.

(43) Siuti, N.; Kelleher, N. L. Decoding protein modifications using top-down mass spectrometry. Nature Methods 2007, 4 (10), 817–821. DOI: 10.1038/nmeth1097.

(44) Kelleher, N. L. Peer reviewed: top-down proteomics. Analytical chemistry 2004, 76 (11), 196 A–203 A. DOI: 10.1021/ac0415657.

(45) Kessner, D.; Chambers, M.; Burke, R.; Agus, D.; Mallick, P. ProteoWizard: open source software for rapid proteomics tools development. Bioinformatics 2008, 24 (21), 2534–2536. DOI: 10.1093/bioinformatics/btn323.

(46) Liu, X.; Inbar, Y.; Dorrestein, P. C.; Wynne, C.; Edwards, N.; Souda, P.; Whitelegge, J. P.; Bafna, V.; Pevzner, P. A. Deconvolution and Database Search of Complex Tandem Mass Spectra of Intact Proteins. Molecular & Cellular Proteomics 2010, 9 (12), 2772–2782. DOI: 10.1074/mcp.M110.002766.

(47) Basharat, A. R.; Zang, Y.; Sun, L.; Liu, X. TopFD: A Proteoform Feature Detection Tool for Top–Down Proteomics. Analytical Chemistry 2023, 95 (21), 8189–8196. DOI: 10.1021/acs.analchem.2c05244.

(48) Horn, D. M.; Zubarev, R. A.; McLafferty, F. W. Automated reduction and interpretation of high resolution electrospray mass spectra of large molecules. Journal of the American Society for Mass Spectrometry 2000, 11 (4), 320–332. DOI: 10.1016/S1044-0305(99)00157-9.

(49) Kou, Q.; Xun, L.; Liu, X. TopPIC: a software tool for top-down mass spectrometry-based proteoform identification and characterization. Bioinformatics 2016, 32 (22), 3495–3497. DOI: 10.1093/bioinformatics/btw398.

(50) Elias, J. E.; Gygi, S. P. Target-decoy search strategy for increased confidence in large-scale protein identifications by mass spectrometry. Nature Methods 2007, 4 (3), 207–214. DOI: 10.1038/nmeth1019.

(51) Choi, I. K.; Liu, X. Top-Down Mass Spectrometry Data Analysis Using TopPIC Suite. In Proteoform Identification: Methods and Protocols, Springer, 2022; pp 83–103.

(52) Zhu, K.; Liu, X. A graph-based approach for proteoform identification and quantification using top-down homogeneous multiplexed tandem mass spectra. BMC Bioinformatics 2018, 19 (Suppl 9), 280. DOI: 10.1186/s12859-018-2273-4.

