## Supplementary_material for "TopDIA: A Software Tool for Top-Down Data-Independent Acquisition Proteomics"

### Table of Content

**Supplementary Figures** ........................................ 3

Supplementary Figure S1. A TD-DIA-MS experiment is divided into cycles, each of which contains an MS1 scan and 20 MS/MS scans. All the cycles in a run are sorted in the increasing order of the retention time and the index of a cycle is its position in the sorted list. 3

### Supplementary Figures

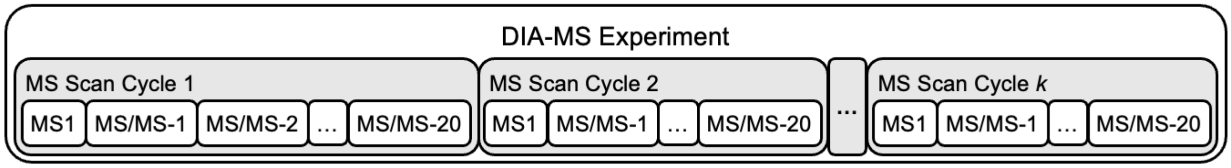

**Supplementary Figure S1.** A TD-DIA-MS experiment is divided into cycles, each of which contains an MS1 scan and 20 MS/MS scans. All the cycles in a run are sorted in the increasing order of the retention time and the index of a cycle is its position in the sorted list.

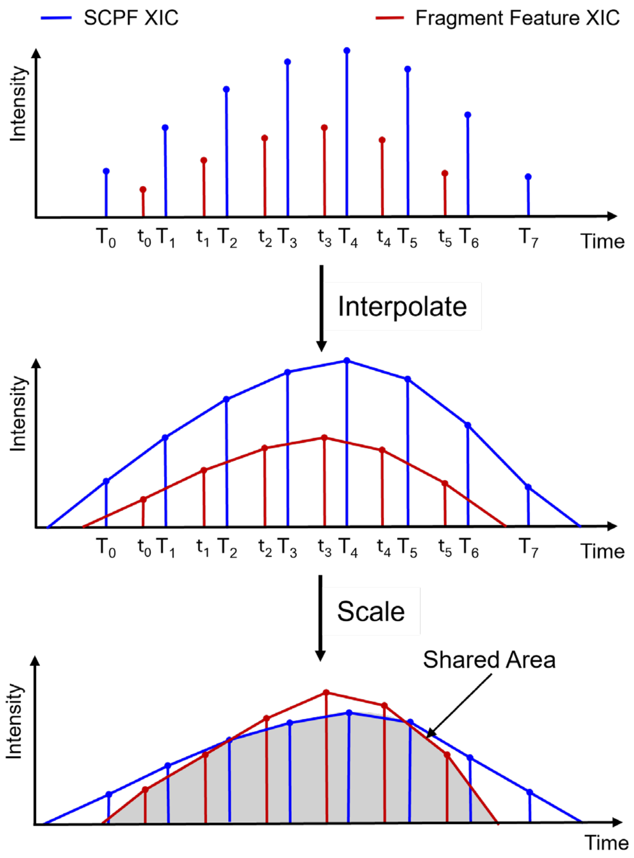

**Supplementary Figure S2.** Extracted ion chromatograms (XICs) of an SCPF and a fragment feature are linear interpolated. Subsequently, the interpolated intensities within the XICs are normalized so that the area under the XIC equals 1. The shared area under the normalized interpolated XICs is reported as the shared XIC of the SCPF and the fragment feature.

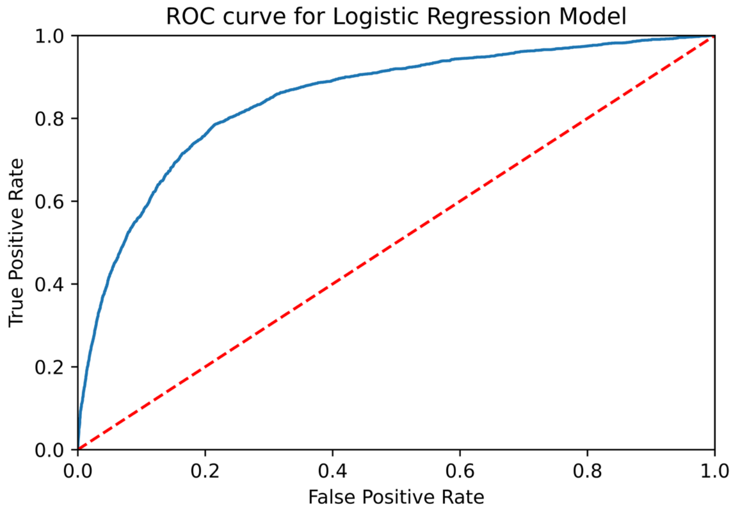

**Supplementary Figure S3.** The receiver operating characteristic (ROC) curve of the logistic regression model on the validation SCPF and fragment feature pairs generated from the DIA-TRAIN data.

### Supplementary Tables

**Supplementary Table S1.** Parameter settings for the modified version of TopFD for extracting proteoform features from TD-DIA-MS data

| **Parameter** | **Value** |
|---|---|
| Maximum charge | 60 |
| Maximum mass | 70,000 Da |
| MS1 signal noise ratio in MS-Deconv | 3.0 |
| *M/z* error tolerance in MS-Deconv | 0.02 |
| Do final filtering in MS-Deconv | True |
| Use EnvCNN score in MS-Deconv | True |
| Use single scan noise level during feature extraction | True |
| Minimum scan number in features | 2 |
| Seed envelope intensity correlation tolerance | 0.5 |
| ECScore cutoff | 0 |

**Supplementary Table S2.** Parameter settings for the modified version of TopFD for extracting fragment features from TD-DIA-MS/MS data

| **Parameter** | **Value** |
|---|---|
| Maximum charge | 60 |
| Maximum mass | 70,000 Da |
| MS/MS signal noise ratio in MS-Deconv | 1.0 |
| Isolation window | 4.0 *m/z* |
| *M/z* error tolerance in MS-Deconv | 0.02 |
| Do final filtering in MS-Deconv | True |
| Use single scan noise level during feature extraction | True |
| Minimum scan number in features | 1 |
| Seed envelope intensity correlation tolerance | 0 |
| ECScore cutoff | 0 |

**Supplementary Table S3.** Parameter settings for TopFD (version 1.7.2)

| **Parameter** | **Value** |
|---|---|
| Maximum charge | 60 |
| Maximum mass | 70,000 Da |
| MS1 signal noise ratio in MS-Deconv | 3.0 |
| MS/MS signal noise ratio in MS-Deconv | 1.0 |
| *M/z* error tolerance in MS-Deconv | 0.02 |
| Do final filtering in MS-Deconv | True |
| Use EnvCNN score in MS-Deconv | True |
| Use single scan noise level during feature extraction | True |
| Isolation window | DDA: 3.0 *m/z*<br>DIA: 4.0 *m/z* |
| Minimum scan number in features | DDA: 3<br>DIA: 2 |
| Seed envelope intensity correlation tolerance | 0.5 |
| ECScore cutoff | 0.5 |
| Additional feature search | False |

**Supplementary Table S4.** Variable PTMs used in TopPIC for protein identification

| PTM | Mass (Da) | Possible amino acid sites |
|---|---|---|
| Methylation | 14.015650 | CKRHDENQ |
| Oxidation | 15.994915 | CPKDNRY |
| Acetylation | 42.010565 | Any |
| Phosphorylation | 79.966331 | Any |

**Supplementary Table S5.** Parameter settings of TopPIC for analyzing TD-DDA-MS and TD-DIA-MS data sets

| Parameter | Value |
|---|---|
| Fragmentation method | File |
| Search type | Target+Decoy |
| N-terminal forms of proteins | NONE, M_ACETYLATION, NME, NME_ACETYLATION |
| Fixed modifications | No |
| Maximum number of variable modifications | 3 |
| Variable modifications | Methylation, Oxidation, Acetylation, and Phosphorylation |
| Spectrum level cutoff type for filtering PrSMs | FDR |
| The cutoff value for filtering PrSMs | 0.01 |
| Spectrum level cutoff type for filtering proteoforms | FDR |
| The cutoff value for filtering proteoforms | 0.01 |
| Error tolerance for precursor and fragment masses | 10 ppm |
| Error tolerance for identifying PrSM clusters | 1.2 Da |
| Maximum number of unexpected mass shifts | 1 |
| Minimum value of the mass shift | -500 Da |
| Maximum value of the mass shift | 500 Da |
| E-values computation | Generating function |
| Use TopFD feature | DDA: True<br>DIA: False |

**Supplementary Table S6.** Comparison of proteoform and protein identifications between DIA-TEST-1 and DDA-TEST-1

| *m*/*z* range | DIA | | DDA | | Shared | |
|---|---|---|---|---|---|---|
| | Proteoforms | Proteins | Proteoforms | Proteins | Proteoforms | Proteins |
| 720-800 | 176 | 84 | 173 | 82 | 122 | 72 |
| 800-880 | 199 | 93 | 209 | 91 | 134 | 78 |
| 880-960 | 205 | 98 | 263 | 109 | 164 | 88 |
| 960-1040 | 202 | 95 | 233 | 101 | 151 | 84 |
| 1040-1120 | 192 | 92 | 190 | 77 | 136 | 68 |
| 1120-1200 | 166 | 78 | 157 | 73 | 113 | 60 |

**Supplementary Table S7.** Comparison of proteoform and protein identifications between DIA-TEST-2 and DDA-TEST-2

| *m/z* range | DIA | | DDA | | Shared | |
|---|---|---|---|---|---|---|
| | Proteoforms | Proteins | Proteoforms | Proteins | Proteoforms | Proteins |
| 720-800 | 151 | 72 | 171 | 74 | 108 | 62 |
| 800-880 | 210 | 94 | 175 | 75 | 126 | 67 |
| 880-960 | 213 | 103 | 227 | 94 | 152 | 81 |
| 960-1040 | 217 | 105 | 225 | 99 | 145 | 80 |
| 1040-1120 | 201 | 97 | 225 | 96 | 148 | 79 |
| 1120-1200 | 134 | 70 | 178 | 79 | 106 | 59 |

**Supplementary Table S8.** Comparison of proteoforms and proteins identified from DIA-TEST-1 by the pseudo spectra and single spectra approaches

| *m/z* range | Pseudo spectra | | Single spectra | | Shared | |
|---|---|---|---|---|---|---|
| | Proteoforms | Proteins | Proteoforms | Proteins | Proteoforms | Proteins |
| 720-800 | 176 | 84 | 167 | 76 | 130 | 72 |
| 800-880 | 199 | 93 | 182 | 82 | 144 | 78 |
| 880-960 | 205 | 98 | 211 | 90 | 154 | 83 |
| 960-1040 | 202 | 95 | 187 | 91 | 153 | 85 |
| 1040-1120 | 192 | 92 | 159 | 73 | 134 | 71 |
| 1120-1200 | 166 | 78 | 155 | 68 | 126 | 63 |

**Supplementary Table S9.** Comparison of proteoform and protein identifications between DIA-TEST-1 and DIA-TEST-2

| *m/z* range | DIA-TEST-1 | | DIA-TEST-2 | | Shared | |
|---|---|---|---|---|---|---|
| | Proteoforms | Proteins | Proteoforms | Proteins | Proteoforms | Proteins |
| 720-800 | 176 | 84 | 151 | 72 | 109 (72.2%) | 64 (88.9%) |
| 800-880 | 199 | 93 | 210 | 94 | 126 (72.0%) | 75 (89.3%) |
| 880-960 | 205 | 98 | 213 | 103 | 145 (70.7%) | 84 (85.7%) |
| 960-1040 | 202 | 95 | 217 | 105 | 143 (70.8%) | 83 (87.4%) |
| 1040-1120 | 192 | 92 | 201 | 97 | 141 (74.2%) | 82 (89.1%) |
| 1120-1200 | 166 | 78 | 134 | 70 | 104 (77.6%) | 63 (90.0%) |

**Supplementary Table S10.** Comparison of proteoform and protein identifications between DDA-TEST-1 and DDA-TEST-2

| *m/z* range | DDA-TEST-1 | | DDA-TEST-2 | | Shared | |
|---|---|---|---|---|---|---|
| | Proteoforms | Proteins | Proteoforms | Proteins | Proteoforms | Proteins |
| 720-800 | 173 | 82 | 171 | 74 | 115 (76.1%) | 66 (94.3%) |
| 800-880 | 209 | 91 | 175 | 75 | 145 (82.8%) | 73 (97.3%) |
| 880-960 | 263 | 109 | 227 | 94 | 161 (78.5%) | 80 (86.9%) |
| 960-1040 | 233 | 101 | 225 | 99 | 160 (79.2%) | 84 (82.3%) |
| 1040-1120 | 190 | 77 | 225 | 96 | 158 (83.1%) | 74 (96.1%) |
| 1120-1200 | 157 | 73 | 178 | 79 | 113 (84.3%) | 62 (93.9%) |
